# Functional characterization of bacterial isolates from dye decolorizing consortia and a step-up metabolic engineering based on NADH-regeneration

**DOI:** 10.1101/2022.04.19.488712

**Authors:** Jagat Rathod, G. Archana

**Author notes:** <u>Corresponding author:</u> G. Archana, Department of Microbiology and Biotechnology Centre, Faculty of Science, The Maharaja Sayajirao University of Baroda, Vadodara-390002, Gujarat, India.

## Abstract

Azo dye decolorizing acclimatized decolorizing consortia are enriched microbial sources of potential azoreductase-efficient bioremediation strains. Here, we characterized eight selected consortial members for their azo decolorization and azoreductase profiling. These efficient dye decolorizing bacterial isolates were affiliated to two major phyla viz. *Firmicute* (genus-*Enterococcus*) and *Proteobacteria* (γ-group). Redox-mediators such as AQDS and AQS were found to significantly increase decolorization except for menadione, and IR functional group signatures highlighted the azo bond reduction and degraded metabolites profiles of each strain. Among isolates, *Enterococcus* sp. L2 was found to be the most effective strain as it could reduce >90mg/L Reactive violet 5R (RV5R) dye in 3h of incubation. Furthermore, strain L2 possesses profound high NADH and NADPH-dependent azoreductase activity which also corroborated with its superior azo decolorization. As per physicochemical parameters, strain L2 showed an optimum decolorization at pH 8, 40 °C and up to 2% w/v salinity. To channelize reducing equivalence (NADH) to further enhance the dye decolorization in NADH-azoreductase efficient *Enterococcus* sp. L2, we augmented an NADH co-factor regeneration system. Using pMGS100, a Gram-positive expression vector a constitutive heterologous expression of *Mycobacterium vaccae* encoded NAD^+^-dependent formate dehydrogenase enhanced NADH pool which led to a significant 3.2 fold increased dye decolorization in *Enterococcus* sp. L2 harboring pMGS100 *fdh* along with a positive effect on growth. Ultimately, an augmentation of formate utilization step could further accelerate azo dye decolorization by fulfilling the co-factor (NADH) requirement of azoreductase along with a growth advantage in the non-model azoreductase-efficient environmentally important strain L2.

## 1. Introduction

Azo dyes are one of the largest classes of dyeing chemicals that account for >70% of the global industrial dye requirement of around 9 million tons (Rawat et al., 2018; Sarvajith et al., 2018; Guo et al., 2020; Routoula and Patwardhan, 2020). Due to their genotoxic and carcinogenic capability, the annual disposal of ~4,500,000 tons of azo dyes and their metabolites are an environmental and financial task (Rawat et al., 2016). For the developing economy, azo compounds and their metabolism, in various biological systems is a top-order agenda of the environment protection and conservation agency which are mainly applied in the textile industry. As per estimates, ~12% of textile-industry associated synthetic azo dyes utilized annually are discharged to wastewater resources. Due to inadequate effluent treatment, these account for ~20% of the total dye pollution to the environment (Saratale et al., 2011). In the last few decades, multiple biological treatments have been attempted using various individual strains and artificial or acclimatized consortia which can bioremediate azo dyes efficiently (Rathod and Archana, 2013; Patel et al., 2016; Rathod et al., 2017; Sreedharan and Bhaskara Rao, 2019; Guo et al., 2021; Samuchiwal et al., 2021), and hunt of new effective strains and their further optimizations remains a challenging task till now.

Azo (-N=N-) bond reduction or decolorization has been the bottle-neck step of the dye degradation pathway, and each strain possesses its unique mode of azo dye decolorization (Sreedharan and Bhaskara Rao, 2019). Multiple mechanisms such as enzymatic, non-specific redox mediator based or direct reduction by reduced metabolites including quinones that can extracellularly reduce azo dye (Stolz, 2001; Chengalroyen and Dabbs, 2013; Rathod et al., 2017)Hong and Gu, 2010).

Enzymatically azoreductase, mono-di oxygenase, peroxidase, laccase, and flavin reductase are the main set of enzymes catalyzing azo decolorization (Chen, 2006; Pandey et al., 2021), out of them azoreductases are recognized critical catalytical component of xenobiotic metabolism and found to be omnipresent in the various biological system (Bafana and Chakrabarti, 2008) (Misal and Gawai, 2018). Azoreductases are also known to possess flavin-dependent quinone reductase activity (Deller et al., 2008; Leelakriangsak, 2013; Suzuki, 2019; Rathod et al., 2022). To catalyze azo cleavage, azoreductases derive reduction potentials from either NADH, NADPH or FADH_2_ (Morrison et al., 2012; Punj and John, 2009). Out of all oxidoreductases, ~80% of enzymes require NADH as a co-factor compared to only up to 10% requiring NADPH (Wu et al., 2013). Therefore, the availability of intracellular NADH to an efficient azoreductase activity remains a competitive metabolic challenge (Rathod et al., 2017). Further, the augmentation of the NADPH-regeneration system is highly complex, and their metabolic requirement of fastidious metabolites (Oeggl et al., 2018), makes it unfavorable for bioremediation application. Previously, we have reported a single gene amended efficient NADH-regeneration system which has been reported to increase azo dye reduction in model bacterial systems. However, their compatibility in non-conventional and environmentally important azo dye reducers has not been studied.

This study aims to characterize azo dye decolorization and azoreductase profiles of unique bacterial strains isolated from our lab enriched acclimatized consortia Rathod and Archana (2013). Among these isolates, we selected the most efficient *Enterococcus* sp. L2 possessing a native NADH-dependent azoreductase activity to further improvise its azo dye decolorization profile. To achieve this, we heterologously overexpress *Mycobacterium vaccae* encoded NAD^+^-dependent formate dehydrogenase in strain L2 to replenish the intracellular NADH pool which is required for efficient azoreductase catalysis. Ultimately, we could accomplish a significant enhancement in azo dye reduction in a non-model and bioremediation points of view important strain.

## 2. Materials and methods

### 2.1. Azo dye decolorization studies

Reactive Violet 5R (RV5R) was used as the model azo dye which consists of a mono azo group linking benzene and a naphthalene ring (Jain et al., 2012; Rathod et al., 2017). RV5R was procured from Meghmani Dyes And Intermediates Ltd, GIDC Vatva, Ahmedabad, India. Methyl red was obtained from HiMedia Laboratories, India. The Bushnell Haas Medium (BHM) [MgSO_4_, 0.2 g/L; K_2_HPO_4_, 1.0 g/L; CaCl_2_, 0.02 g/L; FeCl_3_, 0.05 g/L; NH_4_NO_3_, 1.0 g/L] and various media components used in this study were from HiMedia Laboratories, India. Considering the consortium source of isolate the following media were used, 1) BHM with 0.5% w/v glucose and 0.5% w/v yeast extract as medium A used for isolate ME1; BHM with 0.5% v/v glycerol and 0.5% w/v yeast extract used as medium B for isolate A3; 2% w/v peptone, 0.15% w/v K_2_HPO_4_, 0.15% w/v MgS0_4_, 1% (v/ v) glycerol used as medium C for isolate E2 and K1, and 4) 1.5% w/v Tryptone, 0.5% w/v soya peptone, 0.5% w/v NaCl used as medium D for isolate C1, G1, L1 and L2. Filter-sterilized solution of RV5R and MR dyes was added to obtain its 100 mg/L final concentration in media. To evaluate the NAD^+^-dependent formate dehydrogenase over-expression studies medium with chloramphenicol 10 μg/ ml was used. Dye decolorization at different intervals was monitored by withdrawing aliquots and followed by centrifugation at 14,000 *g* for 10 min to isolate the bacterial cell mass. By measuring the absorbance of the supernatant at maximum wavelength for the Reactive Violet 5R (λ_max_ = 558nm) and Methyl red (λ_max_ = 420 nm) using Spectronic 20D+ (Thermo Scientific) the decolorization percentage was calculated using below equation,

Decolorization (%) = [O.D. at time (t_0_) - O.D. at time (t_1_)] /O.D. at time (t_0_) * 100. Dye decolorization experiments were done in triplicates.

### 2.2. Identification of bacterial isolates by 16S rRNA gene sequencing

For genomic DNA isolation, freshly grown cells were harvested from 2 mL of the culture suspension by centrifugation at 14,000 *g*, 4°C for 10 min. Cell pellet was resuspended in TE25S buffer followed by lysis and purification steps as per standard molecular biology protocols (Sambrook and Russel, 2001). Finally, DNA was dissolved in 50 μl TE buffer (10 mM Tris-Cl pH 8.0, 1 mM EDTA). Using eubacterial universal primers 27F and 1107R, 16S rRNA gene was amplified using PCR (Chaturvedi and Archana, 2012). The PCR product was sequenced using reverse primer (1107R), generating optimum sequence length for the identification (Pillai and Archana, 2008). The sequence data were analyzed using RDP database. MEGA 4.0 was used to construct the phylogenetic tree. Additionally, bacterial identifications were further confirmed using biochemical tests specified by Bergey’s manual (Staley J.R., 2001). Growth analysis was performed by withdrawing cell suspension aliquots at time intervals. To avoid any spectral interference of the residual dye, harvested cells were washed with PBS and growth was measured by taking O.D. at 600nm. Growth experiments were done in triplicates.

### 2.3. Nucleotide sequence accession numbers

All the isolates’ 16S ribosomal RNA sequence GenBank accession number are JQ745287-94.

### 2.4. Azoreductase assay

Azoreductase assay was performed by quantifying the reduction in optical density at λ_max_ of the Reactive violet 5R dye with a Shimadzu UV–visible spectrophotometer at room temperature. The reaction mixture (1.0 ml) contained 25mM potassium phosphate buffer (pH 7.1), 25 μM azo dye, 0.1mM NADH, 10 μM FMN, and a appropriate amount of enzyme. The reaction was initiated by the NADH and quick mixing. One unit (U) of enzyme activity was defined as the amount of enzyme needed to decolorize 1 μmole of azo dye/min/mg of total protein (Chen et al., 2004). Protein concentration was quantified using the Bradford assay (Pierce) and bovine serum albumin (BSA) was used as the standard.

### 2.5. Functional group identification of RV5R degradation products by Fourier Transformed Infrared spectroscopy (FTIR)

Decolorization or degradation products of azo dye by isolates was studied by FTIR analysis. Endpoint metabolites were extracted by an equal volume of ethyl acetate and dried in SpeedVac (Thermo Electron Corporation, Waltham, MA). FTIR Analysis was done by mixing with HPLC grade potassium bromide (KBr) in the ratio of 5:95 and analyzed at mid-IR region (400–4000 cm^−1^) by FTIR using Spectrum GX (PerkinElmer, USA).

### 2.6. *Heterologous expression of NAD^+^-dependent formate dehydrogenase in Enterococcus* sp. *L2*

*Enterococcus* sp. L2 being a Gram-positive isolate, we used pMGS100 *pbacA* including its ribosome-binding site-driven constitutive expression system. The plasmid pMGS100*fdh* was constructed by cloning of coding region (ORF) of myc*fdh* amplified using primers MGS100*fdh*F (5’ ATG GCA AAG GTC CTG TGC GTT CTT TAC G 3’) and Myc*fdh*R (5’ TAT AGG TAC CTT CGG ATC CTC AGA CCG CCTT CTT GA 3’) into NruI site of pMGS100. Clones with correct orientation of *fdh* with constitutive promoter of bacitracin resistance gene (p*bacA*) were screened by BamHI digestion and pcr conformation. *In vitro* handling of DNA molecules for cloning was done utilizing standard protocols (Sambrook and Russel, 2001). The pMGS100 *fdh* was transferred to *Enterococcus* sp. L2 using protoplast electroporation describe by (Dunny et al., 1991).

### 2.7. SDS-PAGE analysis

Cells were harvested and heat-lysed using a boiling water bath for 15 min. A resolving gel (12%) and separating gel were used for SDS-PAGE and as a molecular weight standard protein marker (97, 66, 43, 29, 20, 14kDa) (Merck, India) was used. SDS-PAGE gels were run in 5X Tris-glycine buffer at 70 V for initial 15-20 min and then, at 100 V up to 2h. After electrophoresis, proteins on gels were visualized by staining with 0.25% Coomassie brilliant blue R250 and de-stained according to Sambrook and Russel (2001).

### 2.8. *NAD^+^*-dependent formate dehydrogenase *assay*

To assay NAD^+^-dependent formate dehydrogenase (Fdh) activity, whole-cell lysate was prepared as mentioned for azoreductase activity in sodium phosphate buffer at pH 7.5 along with 0.1M β mercaptoethanol according to Rathod et al. 2017. Using molar extinction coefficient of NADH as 6220 M^−1^ cm^−1^ enzyme units were calculated. One unit (U) of Fdh is defined as the enzyme needed to oxidize 1 μmole formate per minute. Using Bradford method, total protein concentration in cell extracts was measured and bovine serum albumin was utilized as standard.

### 2.9. Intracellular reducing equivalent estimation

Cultures were grown overnight medium containing chloramphenicol 10 μg/ ml. To a re-inoculated freshly grown culture, at the mid-log phase (0.4 O.D.) 1mM IPTG was added which was induced for 6 h along with amendment of 300mM Na-formate. This induced cell culture was centrifuged at 5000 *g* for 10 min and the resulting pellet was washed twice with 0.01M sodium phosphate buffer (pH 7.5). Decanted pellet was resuspended in 1mL 0.01M sodium phosphate buffer (pH 7.5) and sonicated for 3 min by using Sonics VibraCell™, USA. After centrifugation at 14,000 *g*, 4°C for 10 min cell debris were removed and supernatant as cell lysate was used to estimate the reducing equivalents. Using nanophotometer (Implen, GmbH) [NADH] at 340 nm and [Protein concentration] at 280nm was measured, and [NADH]/[Protein] as 340/280 nm ratio was determined.

### 2.10. Statistical analysis

The significant differences among the different treatments were analyzed by one-way analysis of variance (ANOVA) with a pairwise multiple comparison procedure (Fishers LSD). T-test has been performed between treatments of redox mediators to control as well as over-expressing *fdh* transformant to vector control. Sigma Stat 3.5 was used for the statistical analysis.

## 3. Results and Discussion

### 3.1. Azo dye decolorization kinetics of isolates and 16S-rRNA gene-based identification

In our previous study by Rathod and Archana (2013), we have reported the enrichment of twelve acclimatized Reactive violet 5R decolorizing effective consortia from diverse environmental pools. The study also reported a total of 28 isolates from these consortia. These consortia harbored several heterogeneous, active, and profound azo dye decolorizing members which have the potential for efficient azo dye bioremediation. Based on the efficient decolorization properties, out of 28, eight isolates were selected for further taxonomic identification and characterization.

These potential isolates were analyzed for decolorization of complex model dye Reactive violet 5R (RV5R) and Methyl red (mono azo with two benzene rings, MR) in their native growth media of respective consortium. Isolates L2 and ME1 were found to decolorize RV5R more than 90% in 3 h (Figure 1a). Isolate C1 and G1 took up to 30h to decolorize up to 90%. Decolorization of methyl red (mono azo, benzene rings containing dye) was studied for these isolates, resulted 99% decolorization of MR by isolates L2 and ME1 by 6h and isolate C1 decolorized 98% of MR decolorization by C1 isolate in 12 h (Figure 1b), whereas the rest of the isolates took 18h to decolorize MR more than 90%. The chemical structures of the model dyes used in this study are depicted in Figure. 1c, d.

**Figure 1.**
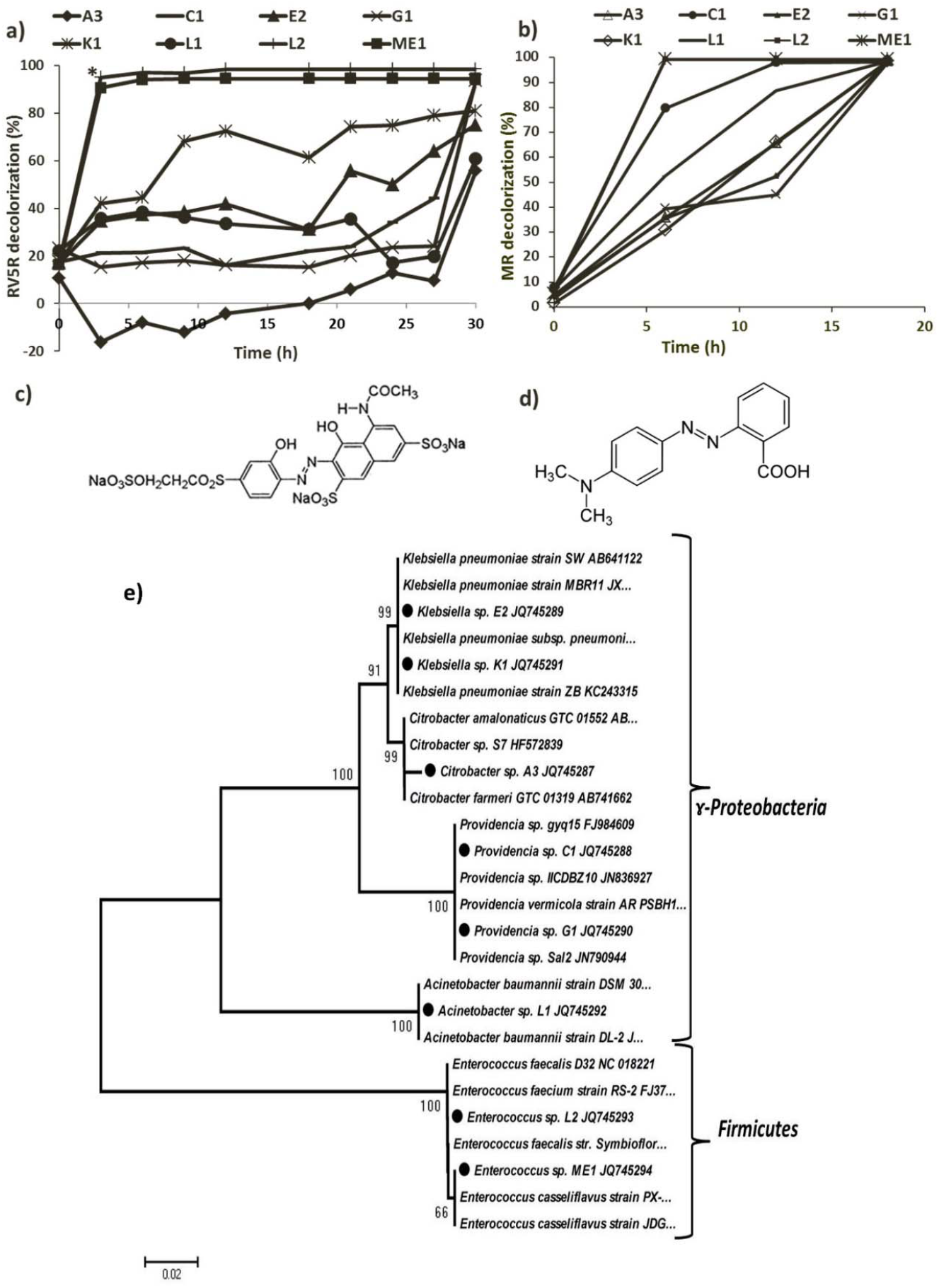
Azo dye decolorization and 16S rRNA gene based phylogenetic analysis of isolates. a) RV5R decolorization; b) Methyl red (MR) decolorization; c) Structure of Reactive violet 5R (RV5R); d) Structure of Methyl red (MR); e) Unrooted phylogenetic tree depicting taxonomic affiliations of the azo dye decolorizing bacterial isolates. Phylogenetic analyses were conducted in MEGA4. RV5R decolorizing isolates of the present study are indicated by dark circles. The percentage of replicate trees in which the associated taxa clustered together in the bootstrap test (1000 replicates) is shown next to the branches. The tree is drawn to scale, with branch lengths in the same units as those of the evolutionary distances used to infer the phylogenetic tree. The evolutionary distances were computed using the Maximum Composite Likelihood method and are in the units of the number of base substitutions per site (scale bar corresponds to 0.2 nucleotide substitution per site). All positions containing gaps and missing data were eliminated from the dataset (Complete deletion option).

Table 1 shows the phylogenetic affiliation of the eight isolates based on 16S rRNA gene sequence. Biochemical key identification results are given in Tables S1-4. The Gram positive were identified as *Enterococcus spp*., whereas six of the gram negatives were found to belong to □-*Proteobacteria*, out of which two belonged to *Providencia* and *Klebsiella spp*., whereas remaining two were similar to *Acinetobacter* and *Citrobacter* genera. Using16S rRNA gene sequence similarities, best matches were selected along with their 16S rRNA sequences from Ribosome data project (http://rdp.cme.msu.edu/) for building phylogenetic tree (Figure 1e). The optimal tree with the sum of branch length was 0.37043799. Further, we correlated the identified member strains with previously reported taxonomic neighbors with dye bioremediation features and their mode of azo dye reduction. *Klebsiella* spp. have been known for micro-aerophilic -aerobic sequential decolorization/degradation process of various textile azo dyes (Franciscon et al., 2009). *Klebsiella spp*. obtained in these studies showed 99% phylogenetic similarity with the reported *Klebsiella* strains showing heavy metal resistance, heavy metals are widely used for the chemical stability of the azo dye and found to be major co-contaminant in the effluents of dye manufacture and application industries. Interestingly, isolate *Klebsiella* sp. E2 showed phylogenetic similarity with copper resistant *Klebsiella pneumoniae* **s**train SW (accession no. AB641122) and *Klebsiella* sp. K1 with nickel resistant *Klebsiella pneumoniae* strain ZB (accession no. KC243315) (Table. 1). Azo-reducing bacteria such as *Shewanella, Citrobacter, Acinetobacter*, *Pseudomonas* have shown to reduce azo dyes with molecular H2, electron donors which includes short-chain fatty acids and redox mediators that are known to profoundly involved in dye decolorization (Hong et al., 2008; Cui et al., 2020). Thus, obtained isolates specifically *Enterococcus* sp. L2 from the current study should be further investigated for their best potentials.

**Table 1.**
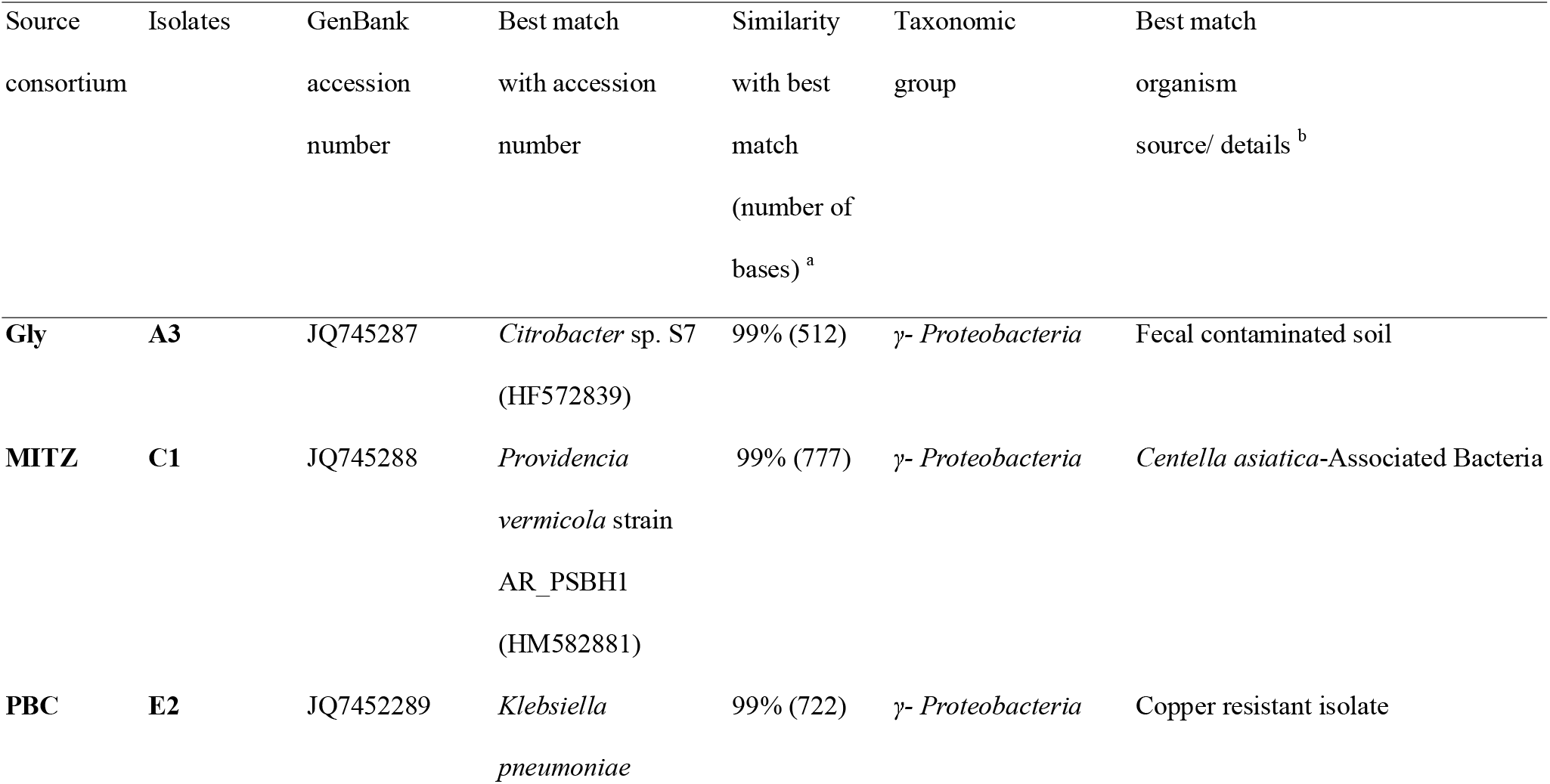
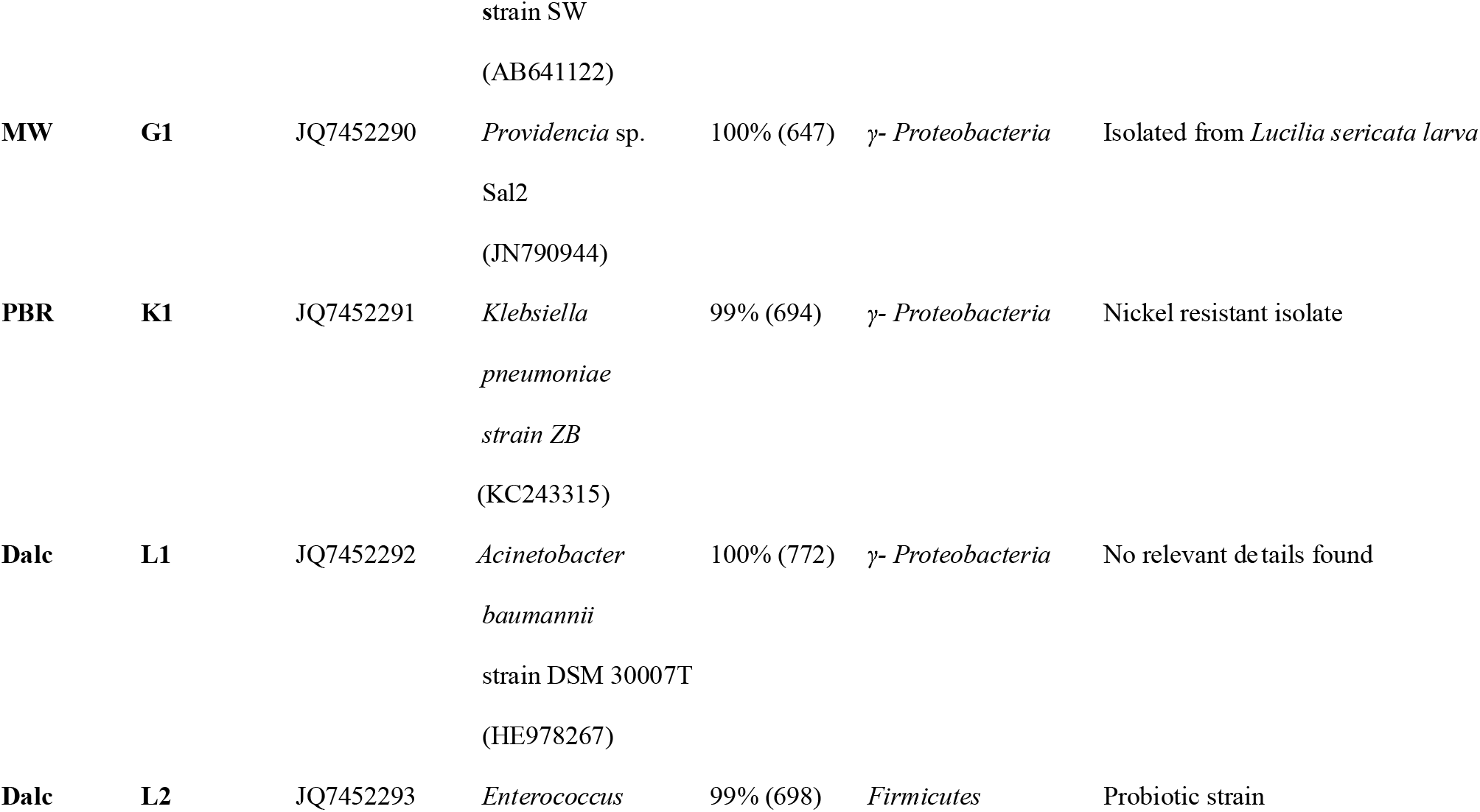
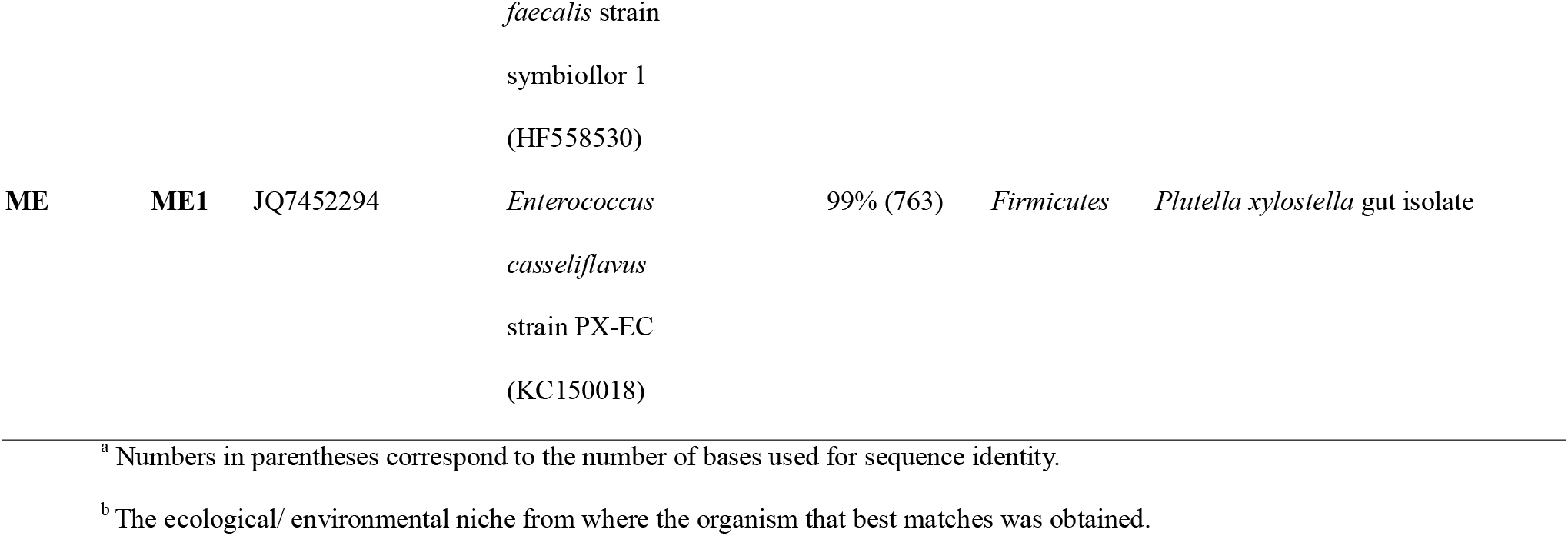
16s rRNA gene sequence based identification of isolates.

| 506 | Source | Isolates | GenBank | Best match | Similarity | Taxonomic | Best match |
| --- | --- | --- | --- | --- | --- | --- | --- |
| 507 | consortium |  | accession | with accession | with best | group | organism |
| 508 |  |  | number | number | match |  | source/ details <sup>b</sup> |
| 509 |  |  |  |  | (number of |  |  |
| 510 |  |  |  |  | bases) <sup>a</sup> |  |  |
| 511 | **Gly** | **A3** | JQ745287 | *Citrobacter* sp. S7 | 99% (512) | *γ- Proteobacteria* | Fecal contaminated soil |
| 512 |  |  |  | (HF572839) |  |  |  |
| 513 | **MITZ** | **C1** | JQ745288 | *Providencia* | 99% (777) | *γ- Proteobacteria* | *Centella asiatica*-Associated Bacteria |
| 514 |  |  |  | *vermicola* strain |  |  |  |
| 515 |  |  |  | AR_PSBH1 |  |  |  |
| 516 |  |  |  | (HM582881) |  |  |  |
| 517 | **PBC** | **E2** | JQ7452289 | *Klebsiella* | 99% (722) | *γ- Proteobacteria* | Copper resistant isolate |
| 518 |  |  |  | *pneumoniae* |  |  |  |
| 519 |  |  |  | strain SW |  |  |  |
| 520 |  |  |  | (AB641122) |  |  |  |
| 521 | **MW** | **G1** | JQ7452290 | *Providencia* sp. | 100% (647) | γ- *Proteobacteria* | Isolated from *Lucilia sericata* larva |
| 522 |  |  |  | Sal2 |  |  |  |
| 523 |  |  |  | (JN790944) |  |  |  |
| 524 | **PBR** | **K1** | JQ7452291 | *Klebsiella* | 99% (694) | γ- *Proteobacteria* | Nickel resistant isolate |
| 525 |  |  |  | *pneumoniae* |  |  |  |
| 526 |  |  |  | strain ZB |  |  |  |
| 527 |  |  |  | (KC243315) |  |  |  |
| 528 | **Dalc** | **L1** | JQ7452292 | *Acinetobacter* | 100% (772) | γ- *Proteobacteria* | No relevant details found |
| 529 |  |  |  | *baumannii* |  |  |  |
| 530 |  |  |  | strain DSM 30007T |  |  |  |
| 531 |  |  |  | (HE978267) |  |  |  |
| 532 | **Dalc** | **L2** | JQ7452293 | *Enterococcus* | 99% (698) | *Firmicutes* | Probiotic strain |
| 533 |  |  |  |  |  |  |  |
| 534 |  |  |  | *faecalis* strain |  |  |  |
| 535 |  |  |  | symbioflor 1 |  |  |  |
| 536 |  |  |  | (HF558530) |  |  |  |
| 537 | **ME** | **ME1** | JQ7452294 | *Enterococcus* | 99% (763) | *Firmicutes* | *Plutella xylostella* gut isolate |
| 538 |  |  |  | *casseliflavus* |  |  |  |
| 539 |  |  |  | strain PX-EC |  |  |  |
| 540 |  |  |  | (KC150018) |  |  |  |

<sup>a</sup> Numbers in parentheses correspond to the number of bases used for sequence identity.
<sup>b</sup> The ecological/ environmental niche from where the organism that best matches was obtained.

### 3.2. Azoreductase profiling of isolates

Among different enzymes catalyzing dye decolorization step, a significant role is contributed by azoreductase in different microbial systems (Liu et al., 2009; Punj and John, 2009; Chen et al., 2010; Husain and Husain, 2012). Azoreductase activity was detected from the isolates, and *Enterococcus* L2 and ME1 had highest NADH- and NADPH-dependent azoreductase activities compared to the rest of the isolates (Table 2). *Enterococcus* L2 and ME1 showed NADH-azoreductase specific activity of 18.73 ± 1.91 and 8.89 ± 1.23, whereas NADPH-azoreductase specific activities were 29.87 ± 2.14 and 15.48 ± 0.57, respectively. Liu et al., (2007) characterized the *azoA* gene from *Enterococcus faecalis* as broad substrate aerobic FMN dependent NADH-azoreductase homodimer of 23kDa subunits. Furthermore, Macwana et al., (2010) characterized *acpD* gene product AzoEf1 from *Enterococcus faecium* as utilized both NADH and NADPH for the reduction of azo dyes. Although, *Enterococcus* sp. L2 has both NADH- and NADPH-dependent azoreductase activities, strengthening the NADH-azoreductase catalysis in strain L2 will be advantageous and physiologically feasible modification as mentioned earlier to optimize azo dye decolorization.

**Table 2.**
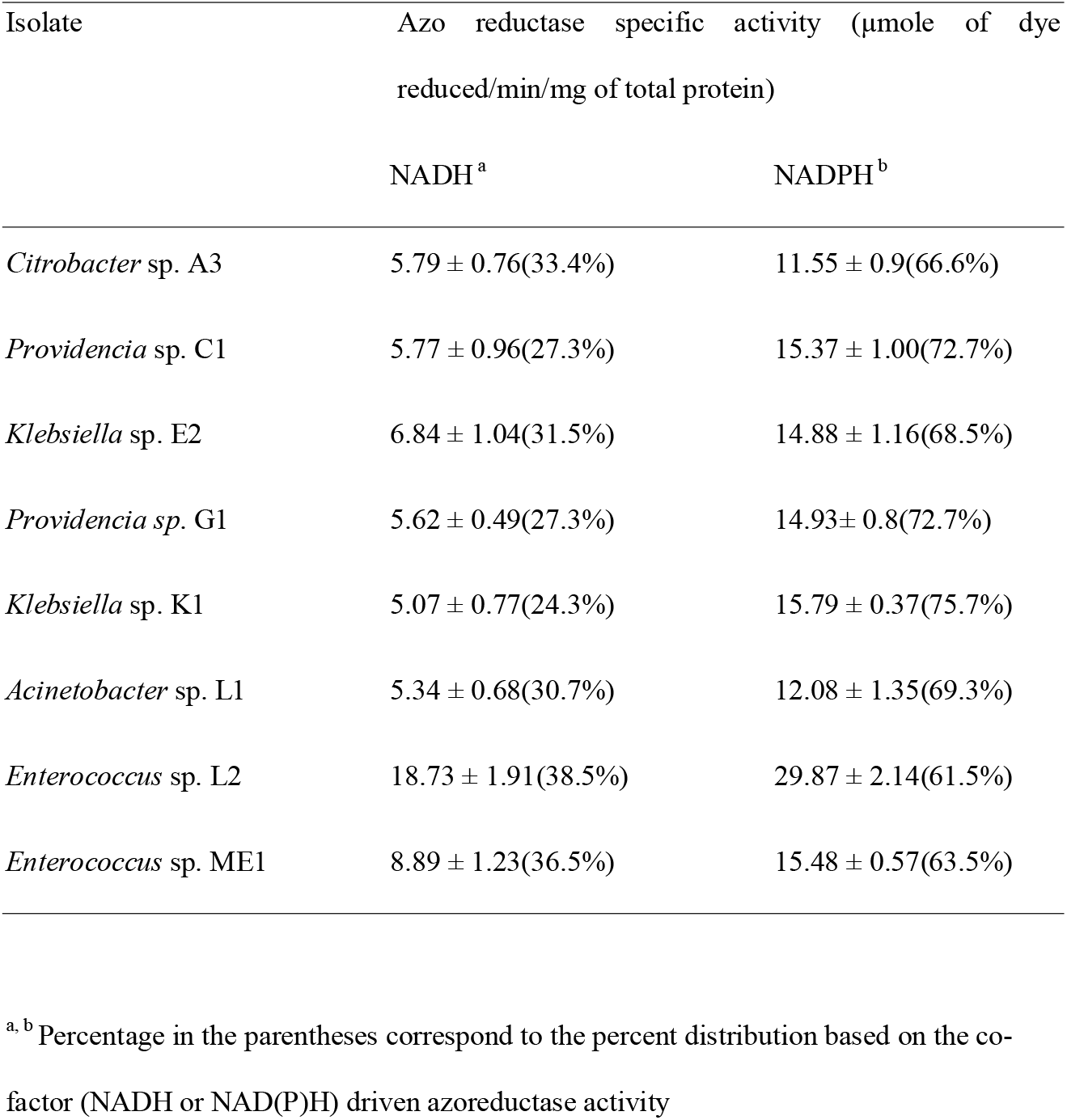
NADH and NADPH dependent azoreductase specific activity of bacterial isolates (Reactive violet 5R as substrate)

### 3.3. Enhancement of RV5R decolorization by preferred redox mediators

Redox mediators involvement in bacterial -N=N-bond reductive cleavage under anaerobic condition have been reported (dos Santos et al., 2003; Van der Zee and Cervantes, 2009; Li et al., 2021), however, their preferences in microbial system and their participation in aerobic conditions for the dye decolorization remains vaguely defined. Flavin enzyme cofactors, such as flavin adenine dinucleotide (FAD), flavin adenine mononucleotide (FMN) and riboflavin, along with other quinone compounds, such as Anthraquinone-2, 6-disulfonate (AQDS), Anthraquinone-2-sulfonate (AQS), and lawsone, are known redox mediators. Most of azoreductases which plays direct role in azo dye decolorization also belong to flavin dependent quinone reductase family, thus physiologically have the ability to accept quinones as substrates (Liu et al., 2008; Rathod et al., 2022). Different concentrations 1.0, 1.5 and 2.0% of Menadione, AQS, AQDS and 1% Lawsone were checked to see the effect of the redox mediators on RV5R decolorization. In the presence of 2.0 mM Menadione the bacterial isolates *Klebsiella spp*. K1 and E2 and *Acinetobacter* sp. L1 had removed approximately double the amount of the dye i.e. 91.82%, 87.89% and 74.46% respectively than the control within 15 h (Figure 2a). In case of increase in the menadione concentration from >1.0 mM enhance decolorization of the RV5R, except *Enterococcus spp. Providencia spp*. showed range specific the positive decolorization effect for menadione which corroborated results by Rau et al., (2002) using menadione. We predict that menadione being electrophilic quinone in nature; imposed oxidative stress on *Citrobacter* sp. A3, *Enterococcus* sp. L2 and *Enterococcus* sp. ME1 which at high concentrations might have led to negative effect decolorization. Significant results were obtained in presence of AQS and AQDS showing ~10 to 20% increase in decolorization at optimum concentration. It was also found that isolates also decrease decolorization beyond optimum concentration of quinones (Figure2b, c). In case of *Klebsiella* strains effect of most of the redox mediators were found to be highly significant, although only *Klebsiella* sp. K1 shown 1.6 fold increases in RV5R decolorization in 1% lawsone (Figure 2d) which is corroborated with the results by Olivo-Alanis et al. (2018). The enhancement mechanism of redox mediators have been elucidated by Zee and Villaverde, (2005), as redox mediators (RMs) accelerate the reaction rate by coupling the microbial oxidation of primary electron donors via shuttling electrons to the acceptor azo dyes (Van der Zee et al., 2003). Yeast extract has demonstrated to improve azo dye decolorization as it can serve as a source of reducing equivalents and electron shuttle which can reduce azo dye (Imran et al., 2016). Hydroxyquinone was also checked for its effect on decolorization, which was found negative (data not shown). Ultimately, a strain-specific effect of RMs was observed on azo dye decolorization as these isolates were equipped by unique set of quinone reductase system which also includes many azoreductases.

**Figure 2.**
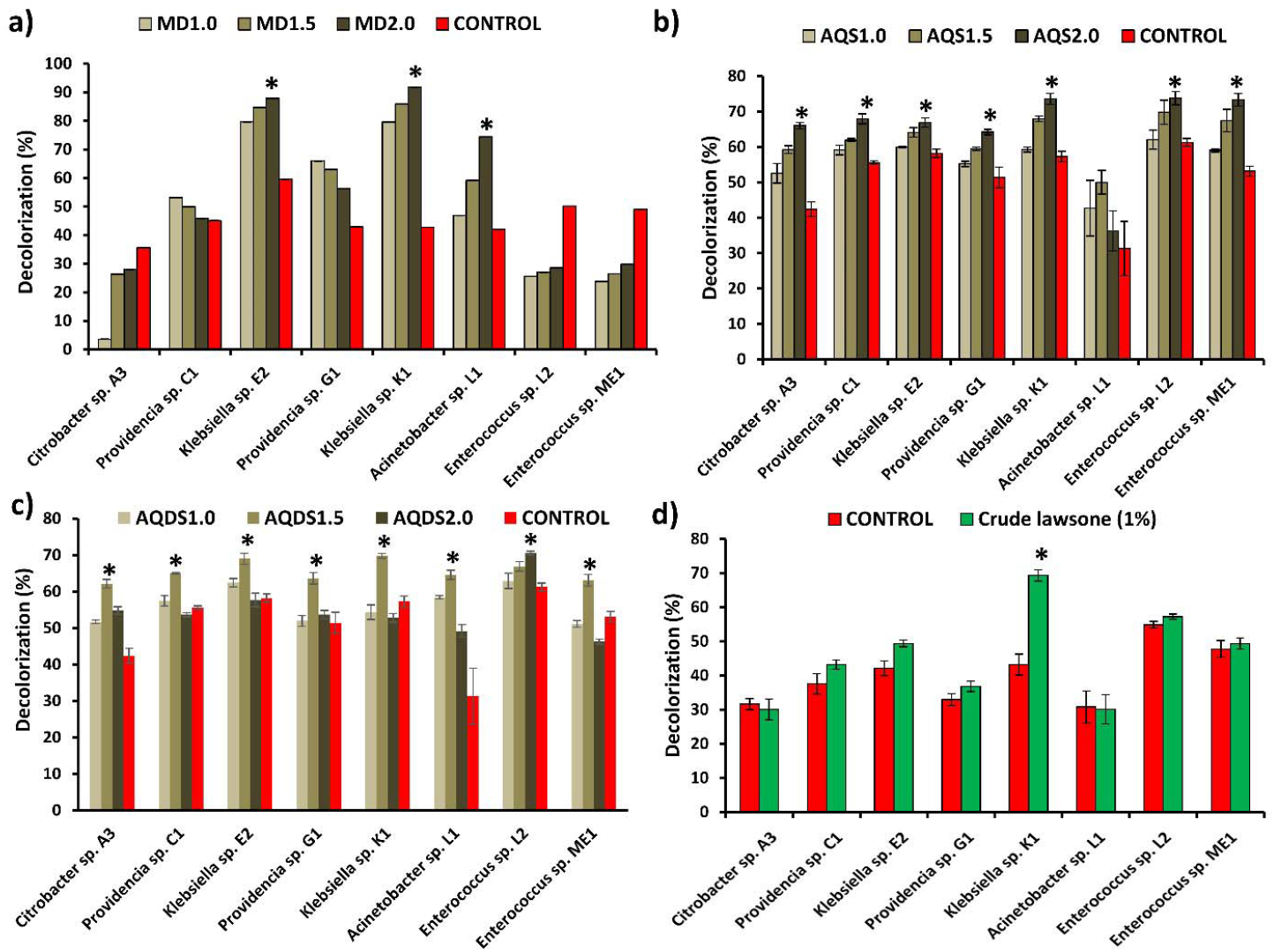
Effect of redox mediators on Reactive violet 5R decolorization. a) Menadione (1, 1.5, 2 mM) b) Anthraquinone-2-sulfonate (AQS) (1, 1.5, 2 mM) c) Anthraquinone-2,6-disulfonate (AQDS) (1, 1.5, 2 mM) d) 1% crude Lawsone. Asterisk sign denotes the statistical significance at *p* < 0.05 for the increase in dye decolorization amended with specific redox mediator at respective concentration compared to control of without amendment of electron mediator.

### 3.4. FTIR analysis of the dye decolorization/degradation end products

Functional groups absorption peaks shifting or dis-appearance in treated samples to control dye sample demonstrates various steps or chemical modifications of the decolorization/degradation process (Jain et al., 2012; Patel et al., 2020). FTIR spectrum form decolorized end product was extracted and compared with the control (RV5R) (Figure S1a-i). Reactive Violet 5R FTIR spectra of as control showed signature peaks for multi-substituted benzene ring along with the peaks at 1,139, 1,185 and 1,547 cm^−1^ which corresponds to two -SO_3_H group, a symmetric SO_2_ and azo bone, respectively (Desai et al. 2009). Azo bond peak at 1547 cm^−1^ was prime signature of a mono-azo reactive azo dye RV5R and loss of this peak in the decolorized extracts of various culture supernatants determined the cleavage of the azo bond (Table 3). FTIR analysis of extracted metabolites of degraded RV5R showed peaks 1630-1680cm^−1^ of primary amines. The peak corresponding to -CN asymmetric stretching at 1048.48 cm^−1^ and -SO3H group 1139.89 and 1185.13 cm^−1^ peak was also disappeared, in all the strains except in *Acinetobacter* sp. L1. Further suggesting that these isolates were capable of removing the sulfonate group from the dye structure and reducing its charge properties, which might enable them to pass through the membrane barrier. The asymmetrical stretching of C–H of alkane (-CH_3_) peak between 2,920-2930 cm^−1^ were observed in degraded metabolites which is corroborated with the findings of asymmetrical C–H stretching in degradation of disperse dye Brown 3REL by *Bacillus* sp. VUS (Dawkar et al. 2008). Thus, these consortial isolates are expected to play a vital and active role in azo dye decolorization and effective bioremediation even as pure culture.

**Table 3.** Fourier transformed infrared spectroscopy (FTIR) analysis of extract end products from the decolorized supernatant from isolates.

| Sample | -N=N-at 1547<br>and<br>1434cm<sup>-1</sup> | Out of<br>plane<br>Aromatic<br>ring C-H<br>bends<br>675-900cm<sup>-1</sup> | Naphthalen<br>e ring at<br>1470cm<sup>-1</sup> | -CN<br>asymmetri<br>c<br>stretching<br>at<br>1048.48c<br>m<sup>-1</sup> | 1139.89,<br>1339.24,<br>1185.13c<br>m<sup>-1</sup> -SO<sub>3</sub>H<br>group | 2920-2930cm<sup>-1</sup><br>asymmetrica<br>l stretching<br>of C-H in<br>CH<sub>3</sub> | 1455-1465cm<sup>-1</sup><br>C-C<br>stretchin<br>g in ring |
| --- | --- | --- | --- | --- | --- | --- | --- |
| Control | + | + | + | + | + | + | shifted |
| A3 | - | - | - | - | - | + | - |
| C1 | - | + | - | - | - | +++ | + |
| E2 | - | + | - | - | - | ++ | + |
| G1 | - | + | - | - | - | ++ | + |
| K1 | - | + | - | - | - | ++ | + |
| L1 | - | + | - | + | - | ++ | + |
| L2 | - | + | - | - | - | + | + |
| ME1 | - | + | - | - | - | ++ | + |

### 3.5. *Effect of* physicochemical parameters such as pH, Temperature, and salinity on dye decolorization by *Enterococcus* sp. L2

The *Enterococcus* sp. L2 was found to decolorize RV5 dye (100mg/L) at an optimum medium pH of 7-8 and temperature 40□C under static conditions (Figure 3a, b). The isolate showed complete decolorization between 35 to 40□C, however sharp decrease in the decolorization was observed above and below this optimum range (Figure 3b). Similarly, Sahasrabudhe et al. (2011) reported *Enterococcus* strain to decolorize Reactive yellow at an optimum pH 5 and temperature for the decolorization at 37▢C. Maximum RV5R decolorization was found to be in the range of 0.5-2% NaCl (Figure 3c), which is the survival and growth range of salinity for *Enterococcus* spp. (Fisher & Phillips, 2009). Recently, similar halo-tolerant and thermophilic bacterial system have been reported for the dye decolorization application (Guo et al., 2021). Interestingly, during *Enterococcus* sp. L2 growth and azo dye decolorization, a significant pH drop was also observed (Figure 3d). Flahaut et. al (1996) reported “flash adaptation” in *E. faecalis*, which makes this bacteria ideal for survival and growth under stress conditions under the bioremediation category. Therefore, *Enterococcus* sp. L2 was selected for additional evaluations.

**Figure 3.**
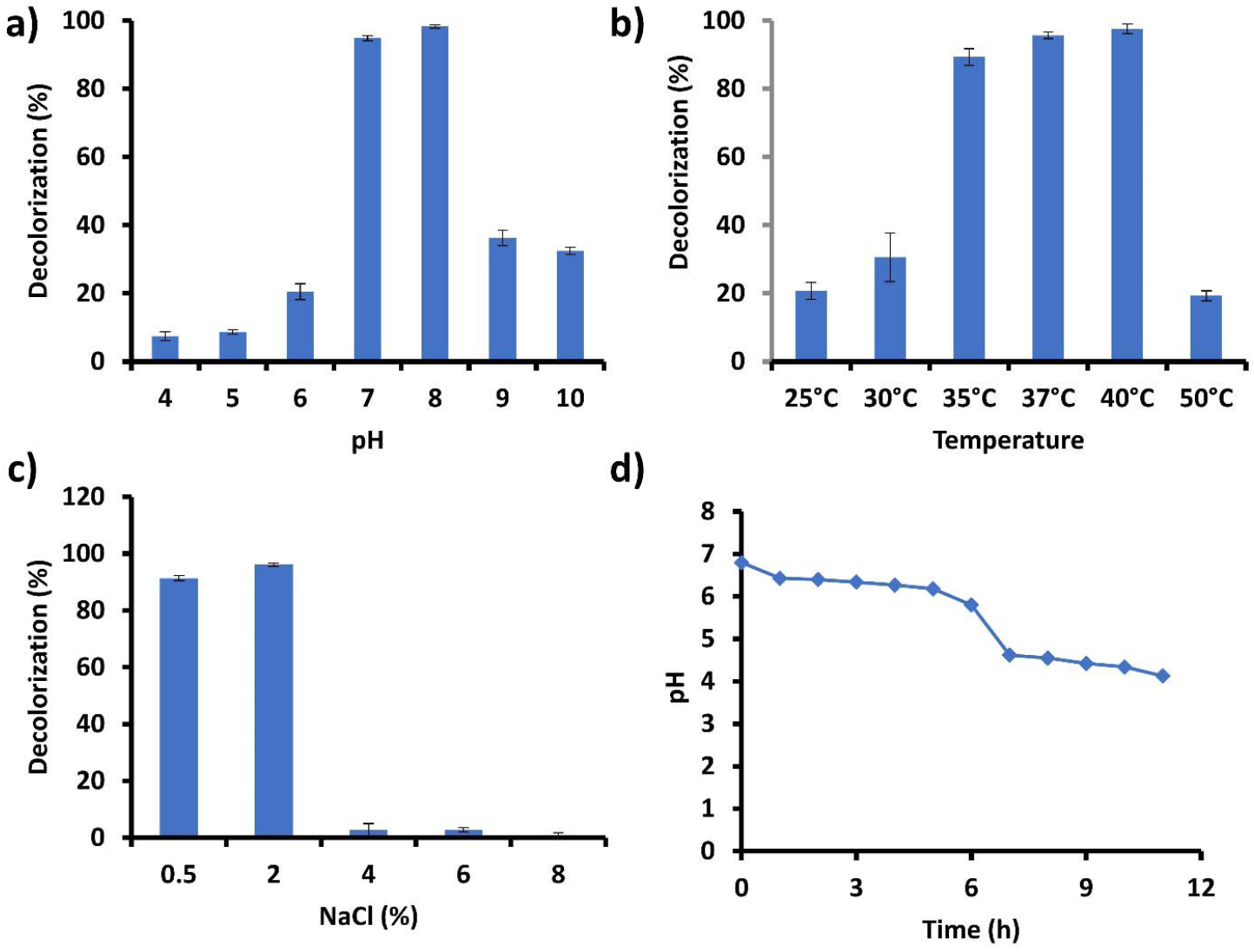
Effect of physicochemical factors on *Enterococcus* sp. L2 Reactive violet 5R decolorization. a) pH; b) Temperature; c) Salinity-NaCl (% w/v) and d) pH-reduction while *Enterococcus* sp. L2 decolorization.

### 3.6. Augmentation of NADH-regeneration systems by heterologous overexpression of NAD^+^-dependent formate dehydrogenase to further enhance decolorization potential of *Enterococcus* sp. L2

The selected isolate, *Enterococcus* sp. L2, was shown to possess NAD(P)H-azoreductase activity, and these reducing equivalences are essential co-factors for azoreductase. While NADH regeneration is physiologically more feasible compared to NADPH_2_ (Oeggl et al., 2018), therefore, we decide to further enhance the NADH to support azoreductase catalysis. To replenish the NADH pool, NAD^+^-dependent formate dehydrogenase was employed which oxidize formate to H2O and CO_2_ while reducing NAD^+^ to NADH. Using pMGS100, a Gram-positive expression vector the *Mycobacterium vaccae* encoded NAD^+^-dependent formate dehydrogenase was heterologously overexpressed by a constitutive *bacA* promoter (Figure 4a). A pMGS100 *fdh* construct was confirmed by BamH1 digestion and a PCR amplification (Figure 4b, c). *Enterococcus* sp. L2 harboring pMGS100*fdh* showed the expected overexpressed protein band of 44kDa (Figure 4d). *Enterococcus* sp. L2 *fdh* transformant showed specific activity of 12.56 U/mg with a fold increase of 6.05 compared to its vector control (Figure 4e). The absorbance ratio of A_340/280nm_ was used as a measure of intracellular NADH concentration relative to the total protein concentration. In medium amended with 300mM Na-formate the average absorbance for A_340/280nm_ ratio for vector control and *fdh* transformant were 0.395 ± 0.009 and 0.455 ± 0.012, respectively. This determined a 1.15 fold NADH increase in *fdh* transformant. Additionally, *Enterococcus* spp. are known to accumulate formate (Leblanc, 2006), and they do not possess formate-hydrogen lyase enzymes or native NAD^+^-dependent formate dehydrogenase activity, therefore, a significant incorporation of final formate oxidation linked to co-factor reductive regeneration. Ultimately, *Enterococcus* sp. L2 *fdh* transformant showed 73.45% decolorization compared to only 22.97% RV5R decolorization by control in 6h, demonstrating a 3.2 fold increase (Figure 4f). This augmentation also led to a significant physiological advantage with positive effect on growth when cell grown with or without supplement of 300mM formate amendment as shown in figure 4g, h. This could be attributed to modified enterococcal system which is now able to utilize formate for the regeneration of NADH when formate was added externally. *Enterococcus* spp. possess pyruvate formate lyase which also naturally produces formate as they could not further utilize it (Leblanc, 2006; Ramsey et al., 2014). Natural accumulation of formate as terminal product of C-metabolism supports the implemented formate dehydrogenase driven NADH-regeneration in *fdh* transformant even when no external formate is added. It is noteworthy that *fdh-based* NADH-regeneration system augmentation in *Enterococcus* sp. L2 could boost its azo dye decolorization and growth.

**Figure 4.**
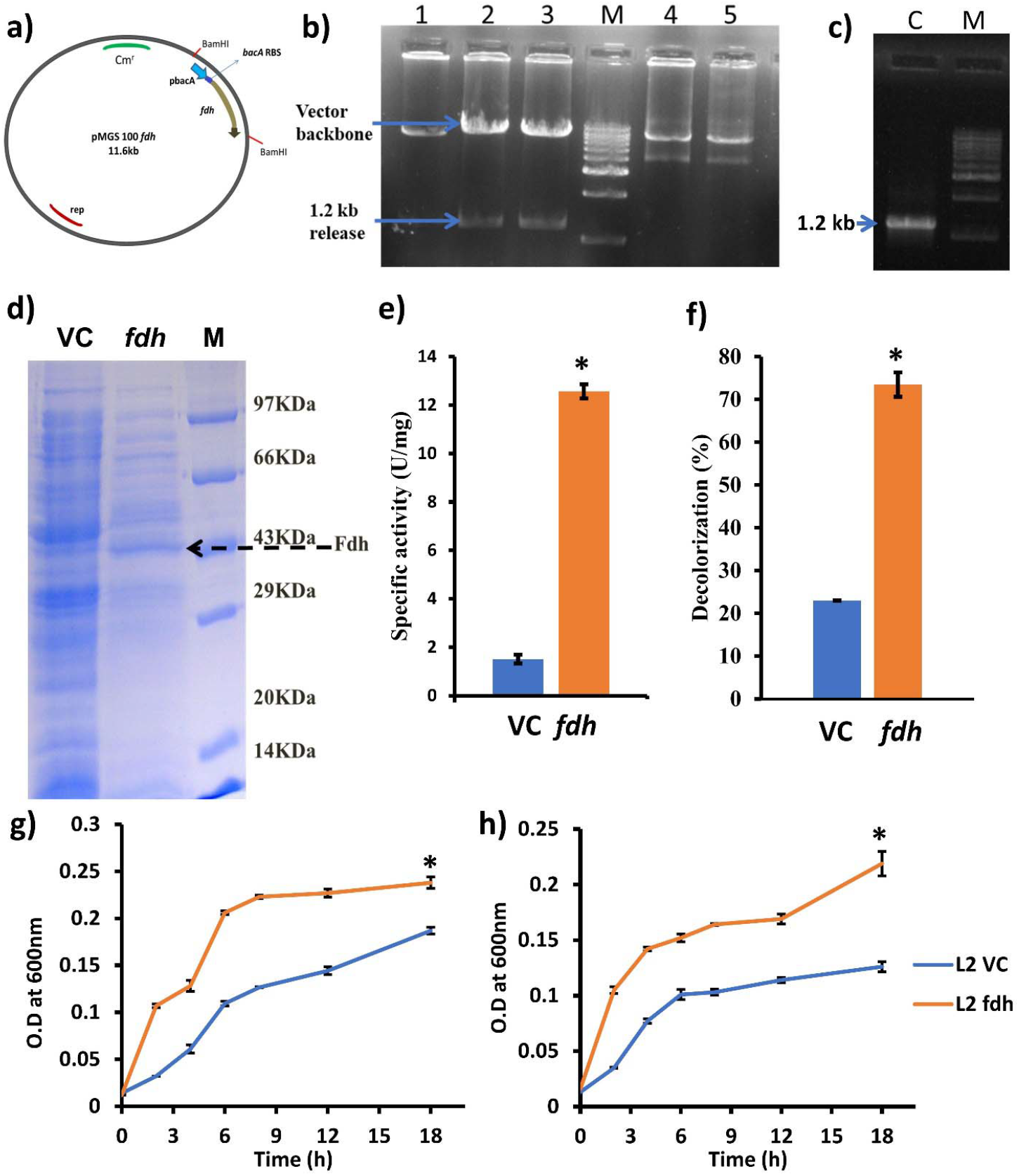
Augmentation of NADH-regeneration systems by heterologous overexpression of NAD^+^-dependent formate dehydrogenase to further enhance decolorization potential of *Enterococcus* sp. L2. a) pMGS100 *fdh* construct map; b) BamH1-digestion confirmation of the *fdh* overexpressing construct and c) PCR confirmation of the *fdh* overexpressing construct; d) Overexpression of 44 kDa protein of *Mycobacterial vaccae* NAD^+^-dependent formate dehydrogenase in *Enterococcus* sp. L2 (VC-vector control, *fdh-fdh-transformant*); e) NAD^+^-dependent formate dehydrogenase activity of *Enterococcus* sp. L2 *fdh* transformant and its vector control; f) Reactive violet 5R decolorization comparison between *Enterococcus* sp. L2 *fdh* transformant and its vector control at 6 h incubation; g-h) Growth comparison between *Enterococcus* sp. L2 *fdh* transformant and its vector control in medium with and without 300mM formate amendment. (Asterisk denotes statistical significance at *p* < 0.01 of increase in Fdh activity and dye decolorization for *fdh* transformant of strain L2 compared to its vector control.)

### 3.7. Potential of NADH-regeneration system in xenobiotic remediation

## 4. Conclusion

Among azo dye decolorizing bacterial isolates from acclimatized consortia, *Enterococcus* sp. L2 was recognized as the most efficient azo dye decolorizer by reducing >90mg/L Reactive violet 5R (RV5R) dye in 3h. A strain-specific preference for redox mediators was demonstrated. A low-cost redox mediator, crude lawsone powder (1%) extract of *Lawsonia inermis* showed positive effect on *Klebsiella* sp. K1’s dye decolorization only. At optimum concentration, AQDS was found to be most preferred redox mediator enhancing dye decolorization in all isolates. It is noteworthy that strain L2 is a NAD(P)H-dependent azoreductase efficient system. Further, strain L2 showed an optimum decolorization at pH 8, 40 °C and up to 2% w/v salinity that were supporting physiochemical features for utilizing strain L2 for biological treatment. NADH-regeneration augmentation in *Enterococcus* sp. L2 by overexpressing NAD^+^-dependent formate dehydrogenase could enhance NADH pool leading to a significant 3.2 fold increased dye decolorization with a positive effect on growth. Ultimately, this study highlighted salient azo dye decolorization traits of strain L2 and its possibility of further optimization by an augmentation of NADH-regeneration system in the non-model azoreductase-efficient environmentally important strain.

## Acknowledgment

Authors acknowledge funding support by the Department of Biotechnology (DBT), Ministry of Science and Technology, New Delhi, India (Project No. BT/PR-6555-BCE/08/424/2005) to GA. JR is thankful to DBT-India for project Junior and Senior Research Fellowships and University Grants Commission, New Delhi, India for Research Fellowship in Sciences for Meritorious Students (UGC-RFSMS).

