## Supplementary Tables and Figure for "Functional characterization of bacterial isolates from dye decolorizing consortia and a step-up metabolic engineering based on NADH-regeneration"

^2^Present address: Department of Earth Sciences, National Cheng Kung University, Tainan, Taiwan-701

*Corresponding author

Corresponding author:

**Supplementary Information**

**Table S1. Biochemical tests for bacterial identification of Gram positive isolates L2 and ME1.**

| **Test** | **L2** | **ME1** | ***Enteococcus***  **standard strain** |
| --- | --- | --- | --- |
| Coagulase | + | + | + |
| Catalase | - | - | - |
| Oxidase | - | - | - |
| Hemolysis | + | + | + |
| Growth in *Streptococcus* *fecalis* broth | + | + | + |

**Table S2. Biochemical tests for bacterial identification of Gram negative isolates E2 and K1.**

| **Test** | **E2** | **K1** | ***Klebsiella***  **standard strain** |
| --- | --- | --- | --- |
| Oxidase | + | + | - |
| Catalase | + | + | + |
| Indole | - | - | +/- |
| Methyl Red | - | - | +/- |
| Voges Proskauer | - | - | +/- |
| Citrate utilization | + | + | +/- |
| Lysine decarboxylase | + | + | +/- |
| Phenylalanine deamination | - | - | - |
| Growth on EMB agar | Green metallic sheen | Green metallic sheen | Green metallic sheen |
| Growth on TSI slant | Slant & butt acidic, Gas formation | Slant & butt acidic, Gas formation | Slant & butt acidic, Gas formation |

**Table S3.** **Biochemical tests for bacterial identification of Gram negative isolates A3 and L1.**

| **Test** | **A3** | ***Citrobacter***  **standard strain** | **L1** | ***Acinetobacter***  **standard strain** |
| --- | --- | --- | --- | --- |
| Oxidase | + | - | - | - |
| Catalase | + | + | + | + |
| Indole test | - | +/- | - | - |
| Methyl Red test | - | + | - | - |
| Voges Proskauer | - | - | - | - |
| Citrate utilization | - | + | + | + |
| Lysine decarboxylase | - | - | - | - |
| Phenylalanine deamination | - | - | - | - |
| Growth on EMB agar | - | - | - | - |
| Growth on TSI slant | - | - | Slant:  alkaline | Slant: alkaline |

**Table S4.** **Biochemical tests for bacterial identification of Gram negative isolates C1 and G1.**

| **Test** | **C1** | **G1** | ***Providencia***  **standard strain** |
| --- | --- | --- | --- |
| Oxidase | - | - | - |
| Catalase | + | + | + |
| Indole | + | + | +/- |
| Methyl Red test | + | + | + |
| Voges Proskauer | - | - | - |
| Citrate utilization | + | + | +/- |
| Lysine decarboxylase | - | - | - |
| Phenylalanine deamination | + | + | + |
| Growth on EMB agar | - | - | - |
| Growth on TSI slant | Slant: alkaline | Slant: Alkaline  Butt: Acidic | Slant: Alkaline  Butt: Acidic |

**Figure S1. FTIR spectra of the parental RV5R azo dye and decolorized end products after the incubation with bacterial isolates.** a) RV5R dye; b) *Citrobacter* sp*.* A3; c) *Providencia* sp*.* C1; d) *Klebsiella* sp*.* E2; e) *Providencia* sp*.* G1; f) *Klebsiella* sp*.* K1; g) *Acinetobacter* sp*.* L1; h) *Enterococcus* sp*.* L2; i) *Enterococcus* sp*.* ME1.
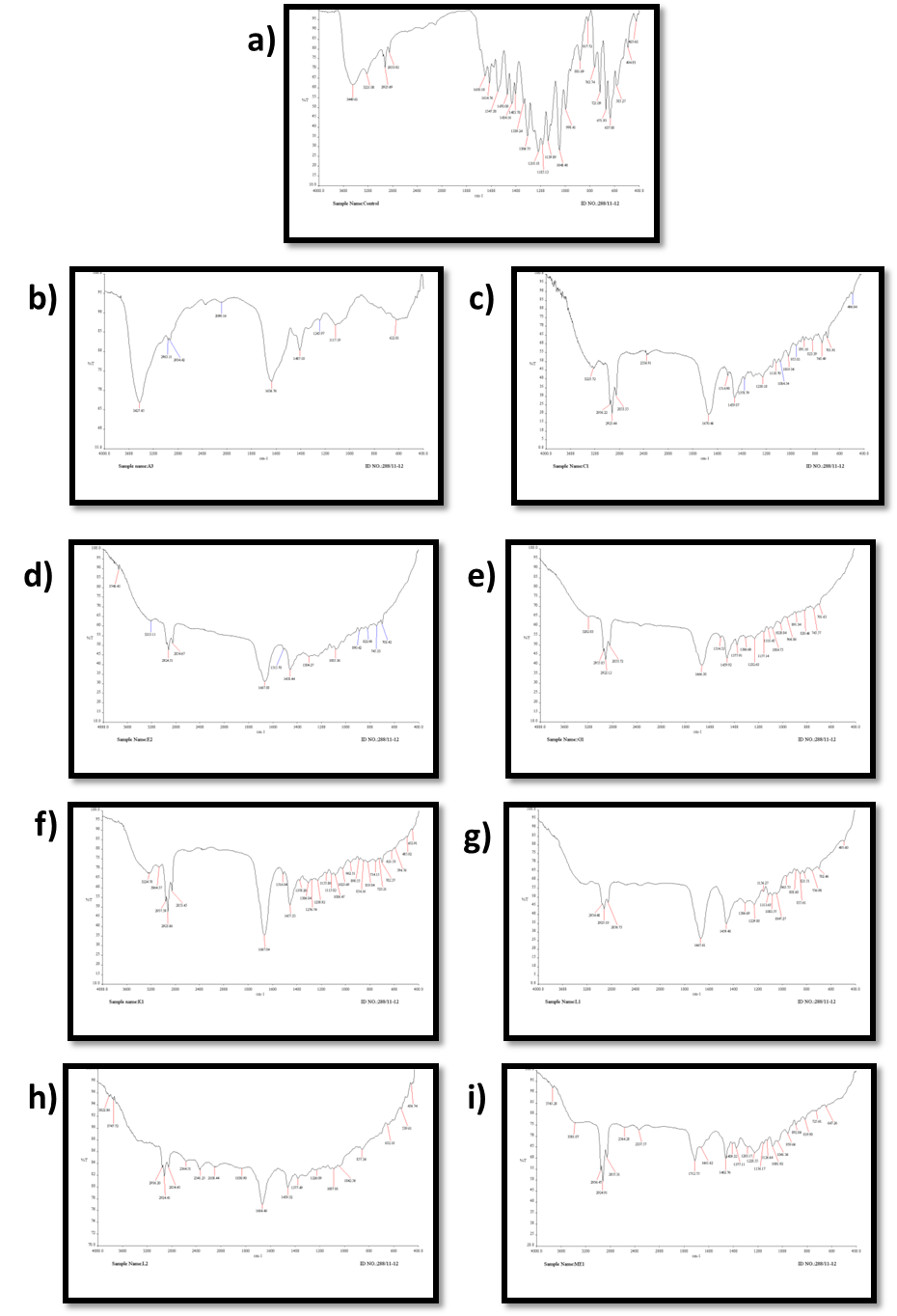
